# Taking advantage of reference-guided assembly in a slowly-evolving lineage: application to Testudo graeca

**DOI:** 10.1101/2024.04.25.591224

**Authors:** Andrea Mira-Jover, Eva Graciá, Andrés Giménez, Uwe Fritz, Roberto Carlos Rodríguez-Caro, Yann Bourgeois

## Abstract

**Background:** Obtaining *de novo* chromosome-level genome assemblies greatly enhances conservation and evolutionary biology studies. For many research teams, long-read sequencing technologies (that produce highly contiguous assemblies) remain unaffordable or unpractical. For the groups that display high synteny conservation, these limitations can be overcome by a reference-guided assembly using a close relative genome. Of chelonians, terrestrial tortoises are considered one of the most endangered taxa, which calls for more genomic resources. Here we make the most of high synteny conservation in chelonians to produce the first chromosome-level genome assembly of genus *Testudo* with one of the most iconic tortoise species in the Mediterranean basin: *T. graeca*.

**Results:** We used high quality, paired-end Illumina sequences to build a reference-guided assembly with the chromosome level assembly of *Gopherus evgoodei.* We reconstructed a 2.29 Gb haploid genome with a scaffold N50 of 107.598 Mb and 5.37% gaps. We sequenced 25998 protein-coding genes, and a 41.2% fraction was determined as repetitive in our assembled genome. Demographic history reconstruction based on the genome revealed two events (population decline and recovery) consistent with previously suggested phylogeographic patterns for the species. This outlines the value of genomes like this for phylogeographic studies.

**Conclusions:** Our results highlight the value of using close relatives to produce *de novo* draft assemblies in species where such resources are unavailable. Our *Testudo graeca* annotated genome paves the way to delve deeper into the species’ evolutionary history and provides a valuable resource to increase making direct conservation efforts on their threatened populations.

## BACKGROUND

Whole genome sequencing (WGS) has become a powerful tool in evolutionary and conservation biology due to the progressive reduction of practical and economical efforts to generate genomic libraries (1). New high-throughput sequencing methods can be used to produce highly contiguous reference genomes for non model or “obscure” organisms (1,2). Long-read DNA sequencing (e.g., Oxford Nanopore Technologies or PacBio) is a promising technique to generate high-quality reference genomes and is established as the future of *de novo* assemblies (3–6). However, long read technologies remain expensive for species with large genomes, and require large amounts of high-molecular-weight DNA to be efficient (6). The involved extraction protocols require fresh or flash-frozen tissues that cannot always be obtained for many laboratories and study systems (3,5,6). Although they are not optimized to produce long contigs, short-read sequencing methods remain more accurate, cheaper and easier to use, even on degraded samples (4–6). Mapping-based and reference-guided assemblies (alignment of contigs/scaffolds to a close relative reference genome) provide a powerful tool to generate contiguous genomes using short reads (4). The long scaffolds obtained from this close relative can be used to anchor the typically short contigs obtained with short reads (4,7). This method is particularly interesting for high conserved syntenic genomes or even with extinct species, where mapping back the reference genome against extant relatives is the only option (4,7–10). Reference-guided assemblies constitute an interesting feedback loop: as more high-quality reference genomes are obtained, there are more chances of a close relative of the species of interest being available (7,9).

Chelonians, the vertebrate group that includes tortoises and turtles, is remarkable for its highly conserved synteny (11,12). The very wide diversity of ecological niches found in chelonians is not reflected in its functional diversity or genome organization, which remains highly conserved across taxa, while nucleotide divergence is particularly reduced and considered a slowly-evolving group (except for mitogenomes) (11–15).

Genetic and genomic resources for chelonians have increased over the past few years. For example, a phylogeny obtained from 98 mitogenomes has contributed to address how ancestral extinctions, niche diversity and biogeography have impacted extant diversity (16). However, microevolutionary processes like gene flow, genomic recombination, introgression or hybridization cannot be extensively addressed using only mitogenomes trees (16). Nuclear reference genomes are also becoming increasingly available. There are currently 36 reference genomes from 13 different families in the NCBI database (Table 1). This source of genomic resources has shed light on speciation events, ancient and recent demographic changes, and are also promising for addressing novel studies, such as genomic determinants of aging, immunology, aridity tolerance or gigantism in chelonians (12–14,17–21). However, the majority of these reference genomes correspond to aquatic species, particularly freshwater turtles (Table 1). The Testudinidae family (land tortoises) is the most threatened family of all chelonians (22), but only counts five annotated genomes, with three from the same genus of American desert tortoises, *Gopherus flavomarginatus*, *G. evgoodei* and *G. agassizii.* The other two include giant tortoises from Galapagos (*Chelonoidis nigra abingdonii*) and Seychelles (*Aldabrachelys gigantea*) (13,14,20) (Table 1). Of them, only three assemblies are annotated at the chromosome level: *G. flavomarginatus*, *G. evgoodei* and *A. gigantea* (Table 1).

**Table 1:**
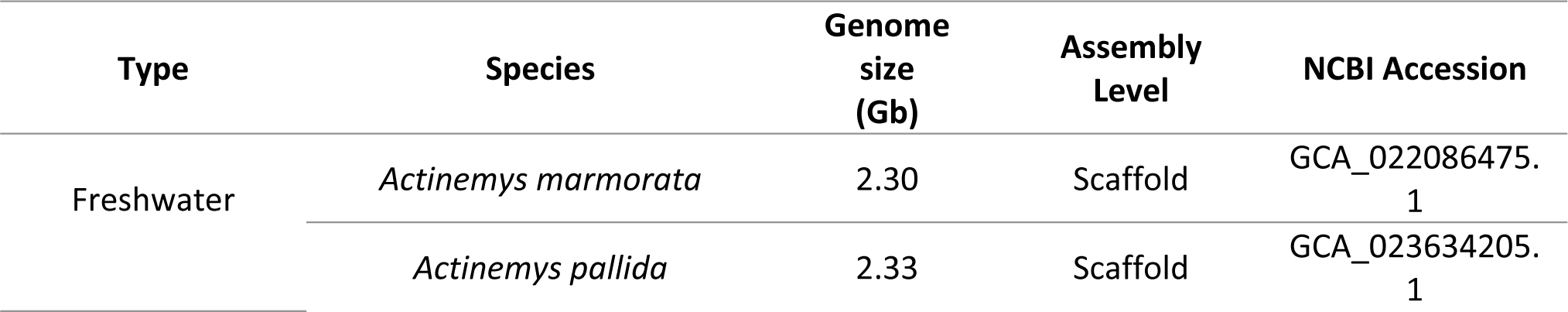
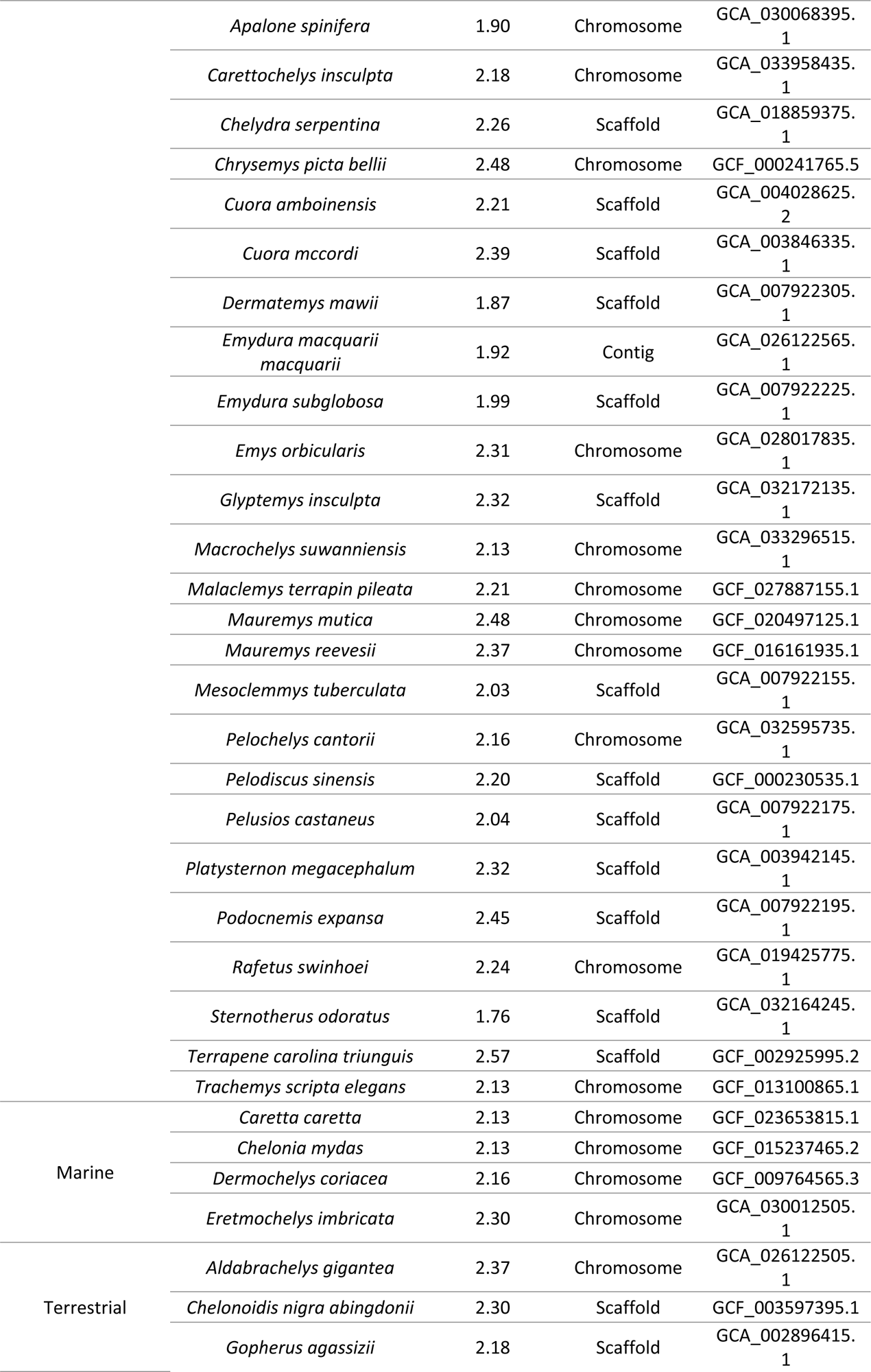
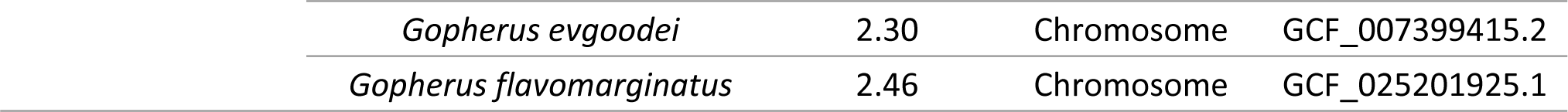
State of the art of chelonians’ reference genome availability. Species are classified and divided into the main chelonians-occupied ecotypes (freshwater, marine or land species). Assembly level represents the highest level for any object in the assembly (i.e., the sequence organization or connection among them). Estimated genome size is represented as Gb for all species. The NCBI database accession number is provided.

| Type | Species | Genome size (Gb) | Assembly Level | NCBI Accession |
| --- | --- | --- | --- | --- |
| Freshwater | *Actinemys marmorata* | 2.30 | Scaffold | GCA_022086475.1 |
|  | *Actinemys pallida* | 2.33 | Scaffold | GCA_023634205.1 |
|  | *Apalone spinifera* | 1.90 | Chromosome | GCA_030068395.1 |
|  | *Carettochelys insculpta* | 2.18 | Chromosome | GCA_033958435.1 |
|  | *Chelydra serpentina* | 2.26 | Scaffold | GCA_018859375.1 |
|  | *Chrysemys picta bellii* | 2.48 | Chromosome | GCF_000241765.5 |
|  | *Cuora amboinensis* | 2.21 | Scaffold | GCA_004028625.2 |
|  | *Cuora mccordi* | 2.39 | Scaffold | GCA_003846335.1 |
|  | *Dermatemys mawii* | 1.87 | Scaffold | GCA_007922305.1 |
|  | *Emydura macquarii macquarii* | 1.92 | Contig | GCA_026122565.1 |
|  | *Emydura subglobosa* | 1.99 | Scaffold | GCA_007922225.1 |
|  | *Emys orbicularis* | 2.31 | Chromosome | GCA_028017835.1 |
|  | *Glyptemys insculpta* | 2.32 | Scaffold | GCA_032172135.1 |
|  | *Macrochelys suwanniensis* | 2.13 | Chromosome | GCA_033296515.1 |
|  | *Malaclemys terrapin pileata* | 2.21 | Chromosome | GCF_027887155.1 |
|  | *Mauremys mutica* | 2.48 | Chromosome | GCF_020497125.1 |
|  | *Mauremys reevesii* | 2.37 | Chromosome | GCF_016161935.1 |
|  | *Mesoclemmys tuberculata* | 2.03 | Scaffold | GCA_007922155.1 |
|  | *Pelochelys cantorii* | 2.16 | Chromosome | GCA_032595735.1 |
|  | *Pelodiscus sinensis* | 2.20 | Scaffold | GCF_000230535.1 |
|  | *Pelusios castaneus* | 2.04 | Scaffold | GCA_007922175.1 |
|  | *Platysternon megacephalum* | 2.32 | Scaffold | GCA_003942145.1 |
|  | *Podocnemis expansa* | 2.45 | Scaffold | GCA_007922195.1 |
|  | *Rafetus swinhoei* | 2.24 | Chromosome | GCA_019425775.1 |
|  | *Sternotherus odoratus* | 1.76 | Scaffold | GCA_032164245.1 |
|  | *Terrapene carolina triunguis* | 2.57 | Scaffold | GCF_002925995.2 |
|  | *Trachemys scripta elegans* | 2.13 | Chromosome | GCF_013100865.1 |
| Marine | *Caretta caretta* | 2.13 | Chromosome | GCF_023653815.1 |
|  | *Chelonia mydas* | 2.13 | Chromosome | GCF_015237465.2 |
|  | *Dermochelys coriacea* | 2.16 | Chromosome | GCF_009764565.3 |
|  | *Eretmochelys imbricata* | 2.30 | Chromosome | GCA_030012505.1 |
| Terrestrial | *Aldabrachelys gigantea* | 2.37 | Chromosome | GCA_026122505.1 |
|  | *Chelonoidis nigra abingdonii* | 2.30 | Scaffold | GCF_003597395.1 |
|  | *Gopherus agassizii* | 2.18 | Scaffold | GCA_002896415.1 |

|  |  |  |  |
| --- | --- | --- | --- |
| *Gopherus evgoodei* | 2.30 | Chromosome | GCF_007399415.2 |
| *Gopherus flavomarginatus* | 2.46 | Chromosome | GCF_025201925.1 |

In the Testudinidae family, the genus *Testudo* comprises five species of Mediterranean terrestrial tortoises (23,24), and three of them are listed as threatened by the IUCN: *T. graeca* and *T. horsfieldii* are considered vulnerable [VU], and *T. kleinmanni* is listed as Critically Endangered [CE]) (22) The spur-thighed tortoise (*T. graeca Linnaeus*, 1758) is the most widespread *Testudo* species in the Western Palearctic and shows an intricate phylogeographic history. Eleven mitochondrial lineages are described for *T. graeca*, and are divided into two different clades. The first, the eastern clade, spans through the Near and Middle East and southeastern Europe, and consists of *T. g. ibera, T. g. terrestris, T. g. buxtoni, T. g. zarudnyi* and *T. g. armeniaca* (25). The second, the western clade, primarily inhabits North Africa, but also includes some isolated populations in southwestern Europe. It is represented by *T. g. graeca; T. g. whitei; T. g. marokkensis; T. g. nabeulensis* and *T. g. cyrenaica* and an additional lineage awaiting a proper description (25). Fossil-calibrated molecular clock analyses based on mitochondrial data suggest that the western clade diverged from its sister taxa, *T. g. armeniaca,* during the Pliocene (7.95-3.48 Mya). Two independent diversification bursts took place during the Mio-Pliocene (8-2 Mya) for the eastern lineages, and during the Pleistocene (1-0.1 Mya) for the subspecies distributed in North Africa (25). In general, the origin of the small populations located in southwestern Europe is recent and located in North Africa (23,25). The species has been historically introduced on Majorca and Sardinia, and in the Doñana National Park (25,26). The exception is the *T. g. whitei* population located in southeastern Spain, with molecular markers indicating a range expansion from North Africa during the Late Pleistocene (20 kya) and subsequent natural expansion in southeastern Spain (23)

To address this lack of genomic resources for this genus, we present the first chromosome-level assembled genome of *T. graeca*. We sequenced it with short-read technology on an Illumina platform to generate a draft assembly, and we used a close relative reference available genome (*G. evgoodei*) to further scaffold and annotate the genome. Our work demonstrates the efficiency of the reference-guided assembly to create accurate *de novo* reference genomes that can serve for future studies by focusing on the complex evolutionary history of *T. graeca* and its conservation.

## MATERIAL AND METHODS

### Sample collection and sequencing

In order to sequence the whole genome of *T. graeca*, we sampled a fresh roadkilled male tortoise from Murcia (southeastern Spain) (S1 Map of sample location). The sample was stored at -4°C. Tissues were extracted under sterile conditions and kept frozen until processing. DNA was extracted from muscle using the E.Z.N.A Tissue DNA kit (Omega Biotek) and eluting it in 100 µL. DNA quantification was performed by a Qubit High Sensivity dsDNA Assay (Thermo Fisher Scientific) at a final concentration of 37.6 ng/µL. Genomic DNA libraries were constructed using the TruSeq Nano DNA kit and quality-checked in the TapeStation D1000 ScreenTape System (Agilent Technologies). Genomic libraries were sequenced on an Illumina NovaSeq with PE150 (paired-end) to obtain a total output of 220 Gb (ca. 100X depth of coverage). Raw FASTQ files were quality-checked using FastQC v0.11.5 (27) (S2 Appendix). All the procedures were carried out by AllGenetics & Biology S.L. following its company protocols.

### Genome assembly

Before starting the assembly, we trimmed all the sequences for remaining adapters and filtered by quality using Trimmomatic 0.39 (28). We discarded reads with a Phred quality score lower than 28 and trimmed reads when quality dropped below 5. We removed the Illumina adapters (TruSeq3-PE) and discarded reads shorter than 40bp. Overlapping reads were merged employing Pear v0.9.11 (default overlap of 10 bp) (29). Sequencing errors were corrected using SOAPec v. 2.0.3 by specifying K-mer size as 27 and the cut-off size as 3 for removing low-frequency k-mers. Assembly was performed using SOAPdenovo2 (version 2.04 release 242) (30) with a range of increasing k-mer values (27, 37, 47, 57, 67, 77, 87, 97, 107). We also tested using k-mer sizes (121 and 127 bp), predicted as being optimal by KmerGenie 1.7051 (31). KmerGenie was also deployed to predict genome size.

The assemblies that employ short reads are generally fragmented and consist in thousands of short contigs. To improve our draft assemblies, we used ntJoin (4) to scaffold our draft assemblies with *G. evgoodei* as a reference given its high-quality chromosome-level assembly with a few unplaced scaffolds. We ran ntJoin with a range of word sizes (100 bp, 250 bp, 500 bp and 1000 bp) and a set of k-mer values (16, 24, 32, 40, 48, 56, 64). Any gaps between contigs were then closed using GapCloser v1.12 (30). To confirm the efficiency of this approach, we examined the completeness of the expected gene in our assembly with BUSCO (BUSCO score v 5.3.0) (32). We tested the continuity and the presence of 5310 Tetraphods shared genes (tetrapoda_0db10) before and after applying ntJoin and always after gap removal with GapCloser). We also compared the quality of the different assemblies by examining N50, NL50 and other statistics using the stats.sh script from the bbmap suite (BBTOOLS 38.18) Quality criteria were assessed according to gaps percentage, the number (NL50) of and the shortest length (N50) longest scaffolds covering half the assembly. Therefore, we retained the assembly with the lowest gaps percentage, the highest N50 and the lower NL50.

### Repeat analysis

In order to identify repeated elements, we used RepeatModeler v2.0.2 (33) to create *de novo* predictions of repetitive sequences and to construct a library of repetitive elements for *T. graeca*. To mask the genome, we combined this *de novo* annotation with an existing consensus of repetitive sequences for Tetrapods using the freely available resources (Dfam) provided with RepeatMasker v4.1.2 (33). We ran the latter program with RMBLAST v2.11.0 to classify and annotate all the repeat families. Then we built a Repeat Landscape to compare *T. graeca* repeat content to other species. We explored the age distribution of Tes by examining the divergence among the different TE families with the calcDivergenceFromAlign.pl script from the RepeatMasker package.

### Gene annotation

Gene finding was performed using BRAKER2 v2.1.6 (34), which incorporates a combination of tools to predict gene coordinates and generates gene structure annotations (34–36). As we do not have access to the RNAseq data for our species, we applied the BREAKER pipeline using the “C” option to incorporate “proteins of any evolutionary distance” into our target species. Because these methods work better with proteins from related species, we combined the protein annotations available for *G. evgoodei, A. gigantea and Gallus gallus* with a set of vertebrate protein data obtained from OrthoDB (tetrapoda_odb10) using DIAMOND (37) to remove any redundant genes between both sources. We ran gene predictions on our masked genome to avoid wrongly annotating Tes as genes. Briefly, the pipeline involves running ProtHint (38) to generate hints of protein prediction by identifying alignments with sequences from close or distant relatives for *T. graeca* in the provided protein database. Annotation is further improved by training AUGUSTUS (34,35,37) on the set of hints to obtain the coordinates and predictions of introns, exons and start/stop codons. We obtained Gene Ontology (GO) terms and gene names for the predicted genes with EggNOG-mapper v2 (39), and by a high-precision search among orthologous groups.

We also transferred the *G. evgoodei* annotation to the *T. graeca* draft genome using Liftoff with default parameters (40).

### Mitogenome reconstruction

We used MitoZ (41) with default parameters on a subset of 10 million pairs of reads to reconstruct the mitogenome of *T. graeca*. Several k-mer values were tested (59, 79, 99, 119, 141). The final assembly was obtained with a k-mer value 141. We aligned our mitogenome reference to other *Testudo* mitogenomes using MAFFT online with default parameters (https://mafft.cbrc.jp/). To further confirm the quality of our sequence, we checked its position in the phylogeny of complete *T. graeca* mitogenomes with a mitogenome from *Testudo marginate* as an outgroup (NCBI accession DQ080047.1). We also employed MAFFT online to run a Neighbor-Joining phylogenetic analysis on the alignment using 100 bootstrap replicates to calculate node support.

### Demographic history inference

Historical changes in the effective population size were inferred with the MSMC2 v2.1.4 software (42). MSMC2 uses a Markov Chain model to estimate the most recent time since coalescence among the haplotypes under recombination. We restricted the analysis to the nine longest scaffolds. To identify poorly mappable regions, we employed GenMap v1.3.0 (43) on the genome assembly. We used BEDTOOLS v2.29.2 (44) to obtain the depth of coverage along the genome (average of 100x). We ran freebayes v1.3.2 (45) to call variants. We masked the regions with a mapability lower than 1 and a depth of coverage below 10X. With VCFtools v0.1.16 (46), we filtered the SNP variants from each chromosome, and excluded the sites with a genotype quality lower than 30 and depth less than 10X or more than 200X. Using the *generate_multihetsep.py* script (provided by msmc-tools, a repository containing utilities for MSMC2, https://github.com/stschiff/msmc-tools), we merged VCF outputs and mask files together to generate the input files for MSMC2. The software was run with default parameters by defining time segmentation as *‘-p 1*2+25*1+1*2+1*3’* and grouping the first and last two-time intervals to force the coalescent rate to remain constant

The coalescence rates estimated by MSMC2 were converted into generations at a mutation rate of 6e-10 bp/year based on the ca. 6% divergence between the *G. evgoodei* and *T. graeca genomes*, which diverged ca. 50 Mya (substitution rate of 3% per lineage over 50 my, or 6e-10 per year) (47). Generation time was estimated at 17.72 years, as in Graciá et al. (2013).

## RESULTS

### Genome sequencing and assembly

For whole genome sequencing, we generated a total of 2 x 913,404,107 high-quality paired-end short Illumina reads with an average sequence length of 151 bp and a GC content of 45%. After adapter and low-quality trimming, we conserved 872,311,739 sequenced reads. *KmerGenie* predicted an optimal k-mer value for the genome *de novo* assembly of 125 bp for an estimated genome size of 2,172,882,866 bp. This estimate is consistent with the sizes obtained for other chelonian genomes assembled at the chromosome level (ranging from 2.13Gb for *Chelonia mydas* and 2.48 Gb for *Chrysemys picta bellii*). Assembling with SOAPdenovo and a k-mer size of 87 produced the assembly with the highest scaffold and contig L50 (5.67 kb and 4.01 kb, respectively; see also Figure S3). This assembly was used for further scaffolding employing ntJoin. A word size of 100 and a k-mer size of 24 resulted in the reference-guided assembly with the highest contig L50 value (3.6 kb) and the smallest gap proportion (13.24%). The proportion of gaps dropped to 5.37% after running GapCloser (Table 2), but the contig L50 rose to 132,837 bp. This scenario suggests that a large proportion of contigs and scaffolds obtained by SOAPdenovo were correctly positioned in relation to one another to ensure efficient gaps filling in the reference-guided assembly (Table 2). The BUSCO complete score rose from 30.3% for the SOAPdenovo assembly to 96.7% after scaffolding with ntJoin and gaps filling, while the proportion of the fragmented and missing genes dropped from 28.8% to 1.1% and 40.9% to 2.2%, respectively (Table 2; Figure 1a).

**Figure 1:**
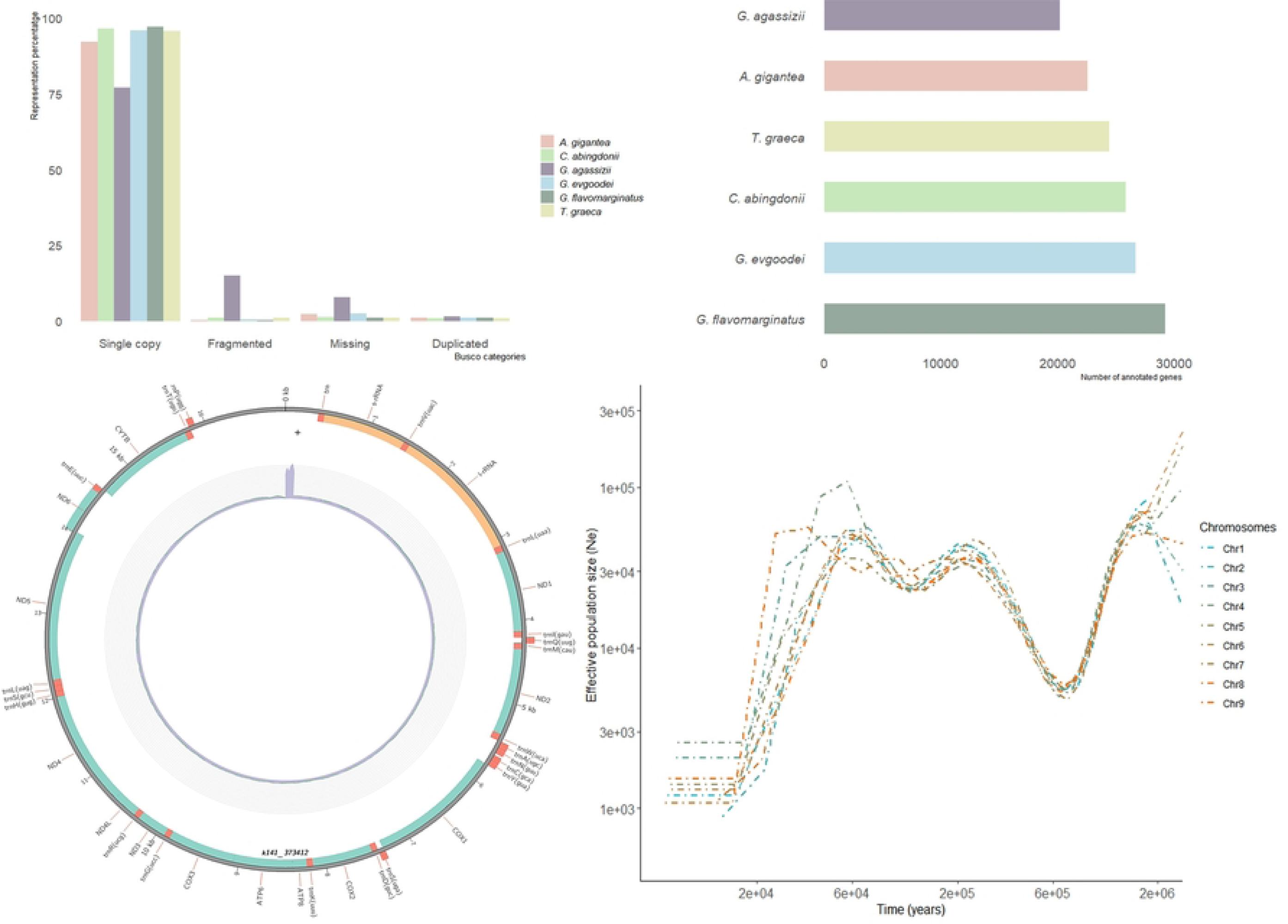
*Representation of the main genome assembly results with the: a) BUSCO completeness comparison with the five assembled Testudineae genomes; b) genes annotated for the five Testudineae genomes and for T. graeca; c) Circos plot of the mitogenome, with the position of the annotated genes and depth of coverage (inner circle); d) demographic reconstruction of the large annotated chromosomes for T. graeca*.

**Table 2:** Comparison of the assembly statics and BUSCO analysis before and after correcting our draft assembly by the ntJoin method.

|  | SOAPdenovo2 assembly<br>(k=87) | Reference-guided assembly<br>after running GapCloser |
| --- | --- | --- |
| **Assembly statics** |  |  |
| *Main genome scaffold (N/L)* | 3895059/2588.160MB | 84,170/2,332.51 MB |
| **Main genome contig (N/L)** | 4236106/2563.699 MB | 112,106/2,207.2 MB |
| **GAPs** | 0.945% | 5.37% |
| **Main genome scaffold N/L50** | 118065/5,676 | 6/130.16 MB |
| **Main genome contig N/L50** | 157154/4,014 | 4,605/132.84 KB |
| **Main genome scaffold N/L90** | 817946/240 | 31/3.92 MB |
| **Main genome contig N/L90** | 1228070/203 | 21,136/13.46 KB |
| **Max scaffold length** | 220,437 | 348.492 MB |
| **Max contig length** | 220,066 | 1.21 MB |
| **Scaffolds > 50KB** | 157 | 144 |
| **% Main genome in scaffolds >50KB** | 0.39% | 92.15% |
| **BUSCO v5.3.0 search** |  |  |
| **Complete and single-copy** | 1561/29.4% | 5082/95.7% |
| **Complete and duplicated** | 49/0.9% | 55/1% |
| **Fragmented** | 1530/28.8% | 56/1.1% |
| **Missing** | 2170/40.9% | 117/2.2% |
| **Total groups searched** | 5310 | 5310 |

### Repeat content

The fraction of repetitive elements identified by RepeatModeler/RepeatMasker was 41.02% for a total of 956,680,633 bp. This falls in line with other related chelonian genomes (Table 3; Figure 2). Long interspersed nuclear elements (LINEs) were the most abundant class of repetitive elements (11.11%), followed by DNA transposons (7.19%), short interspersed nuclear elements (SINEs) (2.06%) and long terminal repeats retrotransposons (LTR-RTs) (3.11%). Unclassified elements accounted for 17.04% of the draft genome. Divergence of repeats from their consensus sequences showed a mode at 7% (S4 Figure), which suggests limited activity of transposable elements (TEs) over recent (> 10 Mya) evolutionary times.

**Figure 2:**
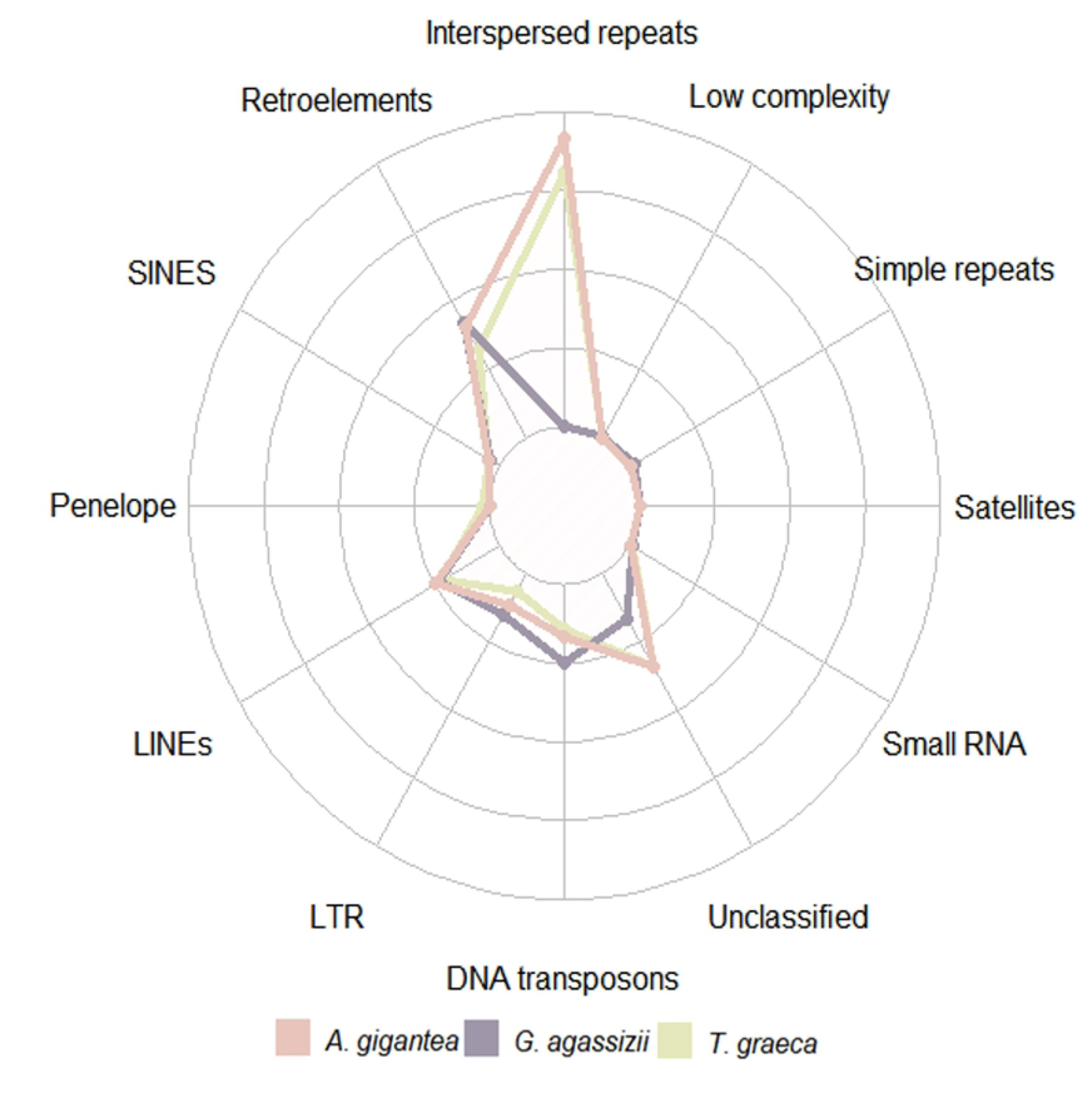
Representation of the different families of repetitive elements found in the Testudo graeca assembly and other closely related available chelonians (Gopherus agassizii and Aldabrachelys gigantea). Testudo graeca and A. gigantea show similar rates of retroelements as DNA transposons over the genome, while the G. agassizii masked region results in a greater presence of retroelements, but fewer DNA transposons. It is important to emphasize that the G. agassizii study only ran RepeatModeler in unmasked regions, while the T. graeca and A. gigantea studies cover the whole genome.

**Table 3:**
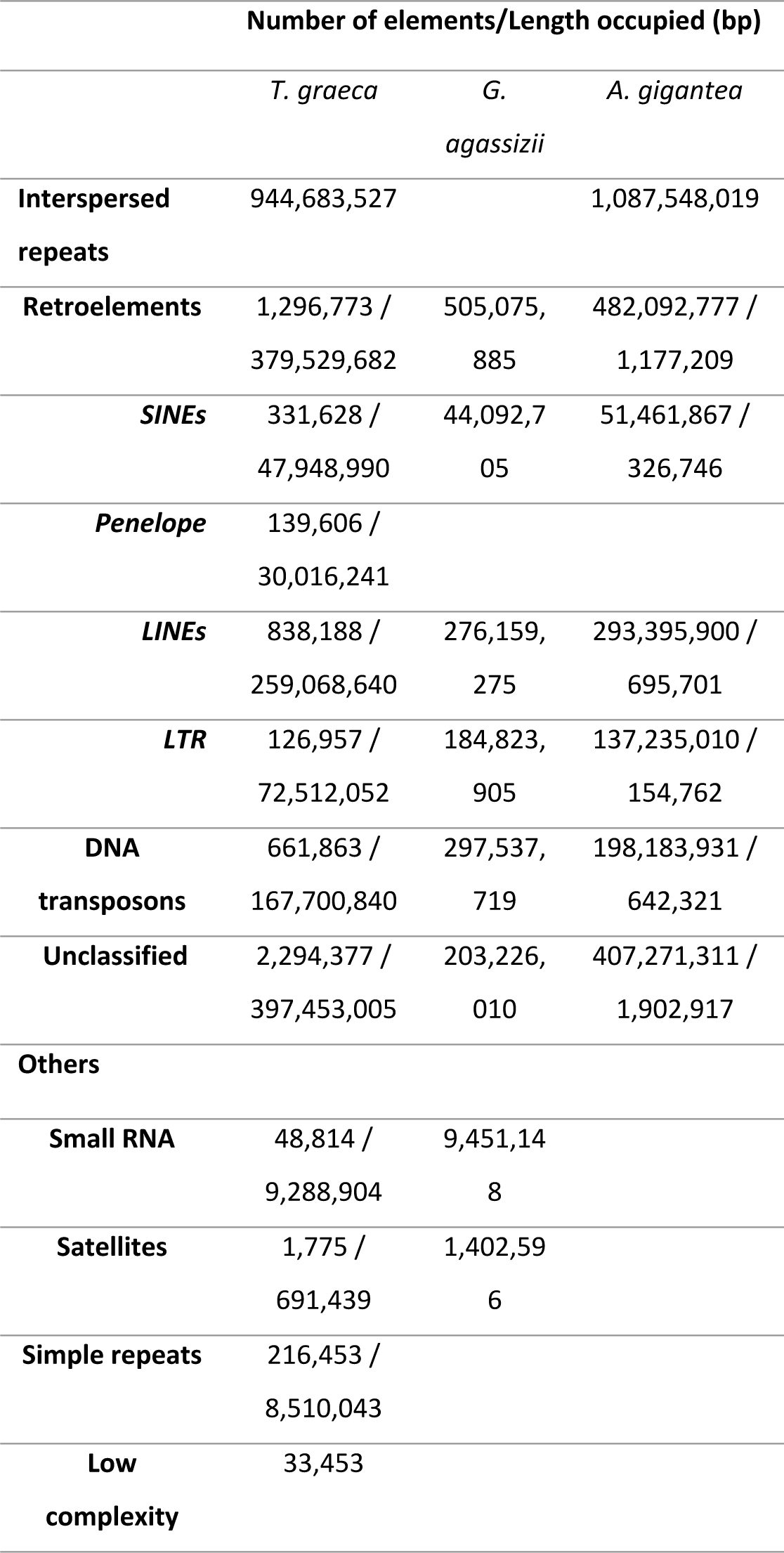
Representation of the different families of repetitive elements found in the Testudo graeca assembly and other closely related available chelonians (Gopherus agassizii and Aldabrachelys gigantea). Testudo graeca and A. gigantea show similar rates of retroelements as DNA transposons over the genome, while the G. agassizii masked region results in a greater presence of retroelements, but fewer DNA transposons. It is important to emphasize that the G. agassizii study only ran RepeatModeler in unmasked regions, while the T. graeca and A. gigantea studies cover the whole genome.

### Gene annotation

The *de novo* annotation of the genome obtained with BRAKER2 using a bank of vertebrate protein sequences recovered 24397 genes with an average length of 6808 bp (Figure 1b). Although the total number of genes recovered falls in line with the number estimated for other Testudinidae, their average length is one order of magnitude shorter of *G. evgoodei* or *A. gigantea* (14). As the *G. evgoodei* annotation benefited from transcriptomic data, we transferred it to our own reference. Of the 19808 coding genes, 462 could not be transferred to the *T. graeca* genome (Table 4). Most (70%) of the genes that were reconstructed *de novo* overlapped with a gene from the *G. evgoodei* annotation, which confirmed that our *de novo* annotation recovered the majority of coding genes, but likely not their full-length sequence.

**Table 4:** Comparison of the de novo annotation with BREAKER and Liftoff of the already existing annotation from the related Gopherus evgoodei.

| Annotation | Number of exons | Number of coding exons | Number of genes | Number of private genes | Average gene length (+/- s.d.) | Average coding exon length (+/- s.d.) |
| --- | --- | --- | --- | --- | --- | --- |
| BRAKER | 129168 | 129168 | 24397 | 7293 | 6808 +/- 10306 | 225 +/- 352 |
| Liftoff from *G. evgoodei* | 324881 | 318841 | 19346 | 2242 | 38670 +/- 67033 | 195 +/- 307 |

### Mitogenome reconstruction and phylogenetic placement

We reconstructed the mitogenome using MitoZ, and obtained a 16928-bp circularized sequence with 37 annotated mitochondrial genes (Figure 1c). Depth of coverage was homogeneous along the sequence (average depth +/- s.d.: 172X +/- 19 when excluding the outlier control region), except for the control region where it peaked at more than 2000X. The comparison to the other three complete *Testudo graeca* mitogenomes (NCBI accession numbers: DQ080047.1, DQ080049.1, DQ080050.1) confirmed the completeness of our assembly outside the control region, and the whole sequence was fully aligned to the three other references ranging from 96.2% to 98.7%. Our mitogenome was shorter than the other references (lengths ranging from 17674 to 19278bp) due to a shorter control region. This region could not be fully assembled due to its repetitiveness and the relatively short length of our reads and inserts. A Neighbor-Joining phylogenetic analysis places our reference close to the Tunisian samples with high support, but more distant from the Turkish sample. All this is consistent with expectations given the species’ biogeography (Figure S5).

### Demographic reconstruction

To evaluate the interest of *T graeca*’s reference genome for demographic analyses, we reconstructed its past demographic history using MSMC2 (Figure 1d). We estimated effective an population size (Ne) to have revolved around 30000 individuals over the last 3 million years, with two population decline events: the first one around 1 Mya, when Ne declined to ca. 6000 individuals; the second decline more recently occurred at 40-20 kya, with Ne declining to ca. 1000 individuals. Of these declines, we observed a significant recovery in Ne, approximately by one order of magnitude, during the period between 600-200 kya.

## DISCUSSION

By means of Illumina NovaSeq PE150 sequencing, we generated the first high-quality draft genome assembly for *Testudo graeca,* including its mitogenome. The reference-guided assembly notably increased sequence contiguity and facilitated annotation. This illustrates the efficiency of the reference-guide assembly for chromosome-level scaffolding and gene annotation by providing a resource for comparing genome organization and diversity within and across clades (4,48). Taking *G. evgoodei* as a reference drastically reduced the presence of any “fragmented” and “missing genes”. BUSCO completeness scored favorably with other chelonians (Figure 1a), but it should be noted that Çilingir et al. (2022) conducted a BUSCO analysis using OrthoDB v10 datasets from phylum (vertebrata_odb10) and class (sauropsida_odb10) instead of all the tetrapods.

BRAKER2 gene prediction estimated a similar number of genes to other Testudines (Figure 1a, b). However, lack of RNA-seq data prevented us from obtaining full-length transcripts and genes. By making the most of the high contiguity of our reference, and combined with conserved synteny and high identity with *G. evgoodei*, we were able to transfer the annotation of the latter to the *T. graeca*’s genome (Table 4).

As *T. graeca* shows accurate differences between population and lineages, the genome herein produced is valuable for further population genomics studies. Using reference genomes from distantly related species can negatively impact SNP calling by underestimating the number of variants or biasing heterozygote calling (49). This effect is significant in turtles and tortoises (12) and, thus, obtaining a reference from the same species ensures future accurate genotyping to avoid a bias in the analyses. Demographic history reconstructions can answer biological questions and trace evolutionary signals to shape the current distribution range and the population genetic status of species (42,50). The population size dynamics herein inferred aligns well with the past population changes proposed in previous studies that used mtDNA or STR, and to show consistency, even at the population level (23,25). The more ancient population decline is compatible with the rapid radiation suggested for the North African lineages during the Pleistocene (25). Today these subspecies show a clear niche differentiation in North Africa, particularly in relation to climate variables like rainfall (51). The subsequent population recovery aligns with the diversification of *T. g. whitei*, and has been estimated to have occurred between 850 and 170 kya (number for Graciá et al. 2017). For this lineage, it has been suggested that it is confined to several refuge areas during the Last Glacial Maximum, from which it subsequently expanded (51), and a similar pattern can be anticipated during other glacial maxima. Repeated contractions and expansions may have greatly contributed to lineage diversification during the Pleistocene. Finally, the more recent decline is consistent with the bottleneck linked with the species’ arrival in southeastern Spain, estimated to have occurred some 20 kya years ago (23,25). As inferred for *T. g. whitei* in our demographic reconstruction, the currently available reference genome for the species would increase knowledge of past population dynamics and other lineages’ demographic history.

Our repetitive element analysis showed that half the genome was made of TEs, which is a similar proportion to other relatives, such as *C. mydas*, *C. picta bellii* or *Gopherus* spp., although assemblies are less than the estimates for *A gigantea* (46.7%) or *T. s. elegans* (45%) (Table 2, Figure 2). The Interspersed Repeat Landscape suggests very low recent transposition given the observed age distribution of TEs. Recent TEs activity appears unlikely, and is possibly biased due to the difficulty of assembling highly repeated regions, and using reference-guided ones. However, this is unlikely given the Kmergenie estimates, which are consistent with the reconstructed genome length and consistent estimates with a c-value.

## CONCLUSION

In this study, we report the first reference genome for the *Testudo* genus by adding to the “toolkit” of genomic resources for terrestrial tortoises. Given the group’s shared synteny, we made the best of the high-quality assemblies of relative Testudines like *G. evgoodei* to scaffold and annotate a chromosome-level assembly genome from short-read sequences. Thanks to this approach, we avoided the higher costs and sample management issues of long-read techniques, while making the most of the low error rate and the cost effectiveness of short read sequencing.

This newly generated reference genome will be useful for answering open questions about the evolutionary history and conservation of the *T. graeca* complex, and possibly of other *Testudo* species.

## ACKNOWLEDGMENTS

This work was supported by Project PID2019-105682RA-I00 and TED2021-130381B-I00, funded by the Spanish Ministry of Science and Innovation (State Research Agency) (MICIU/AEI/10.13039/501100011033), the last also with the support of the European Union “NextGenerationEU”/PRTR”. We thank AllGenetics SL sequencing services, Portsmouth University for computational resources and to all the Ecology Area of Miguel Hernández University (specially to Paco Botella), Serbal, Ecologistas en Acción de Murcia and Andalucia and Murcia Region governments for their field support. Finally, we acknowledge all the projects whom provide new and high-quality reference genomes of non-model organisms.

## Authors contribution

***Andrea Mira-Jover***: Formal analysis; Visualization; Writing-Original Draft; Writing-Review and Editing

***Eva Graciá***: Conceptualization; Founding Acquisition; Supervision; Resources; Visualization; Writing-Review and Editing

***Andrés Giménez***: Founding Acquisition; Supervision; Resources; Visualization; Writing-Review and Editing

***Uwe Fritz***: Validation; Visualization; Writing-Review and Editing

***Roberto Carlos Rodriguez-Caro***: Resources; Validation; Visualization; Writing-Review and Editing

***Yann Bourgeois***: Formal analysis; Conceptualization; Resources; Supervision; Visualization; Writing-Original Draft; Writing Review and Editing.

